# Sex-dimorphic and age-dependent organization of 24 hour gene expression rhythms in human

**DOI:** 10.1101/2022.06.13.494714

**Authors:** Lorenzo Talamanca, Cédric Gobet, Felix Naef

## Abstract

The circadian clock modulates most of human physiology. However, the organization of tissue-specific gene expression rhythms is still poorly known, as well as age and sex dependencies. We leveraged the Genotype-Tissue Expression project (GTEx) using a novel algorithm to assign a unique circadian phase to 914 individuals and transfer time information from stronger to weaker clocks. These donor internal phases allowed us to identify and compare programs of rhythmic gene expression in 46 tissues. Clock transcripts showed highly conserved phase and amplitude relationships across tissues, and were tightly synchronized across the body. Tissue rhythmic gene expression programs differed in breadth, covering global and tissue-specific functions, including metabolic pathways and systemic responses such as heat shock. The circadian clock structure and amplitude was indistinguishable across sexes and age groups. However, overall gene expression rhythms were highly sex-dimorphic and more sustained in females. Moreover, rhythmic programs dampened with age across the body. Together, our stratified analysis unveiled a rich organization of sex- and age-specific circadian gene expression rhythms in humans.

**One sentence summary:** Circadian phase inference of 914 GTEx donors reveals sex- and age-dependent rhythmic gene expression programs across 46 human tissues.

## Introduction

The circadian clock is the evolutionary adaptation of life to the 24h periodicity of earth rotation. The clock synchronizes internal body rhythms in behavior and physiology with external cues, such as 24h environmental or societal Zeitgebers or feeding rhythms [1–3]. Perturbations of the clock, as caused by sleep disruption and shift work can lead to metabolic pathologies [4–7]. sexual dimorphism and its implications in human physiology are well studied [8], recently also in terms of gene expression differences across the body [9]. Many complex phenotypes, including diseases, exhibit sex-differentiated characteristics [10]. However, interactions between sexual dimorphism and circadian rhythms in humans are poorly known [11]. Likewise, the effects of aging on human physiology are well studied [12], notably in cardiovascular tissues [13]. Circadian dysfunction has been observed in brain disorders often emerging at later stages of life [14]. The interplay between circadian oscillations and aging processes are still poorly known and mainly studied in mice [15,16].

Here we combined GTEx transcriptomes with a novel internal phase assignment algorithm that significantly extends existing approaches [17–19]. and allowed us to obtain a whole organism view of 24h gene expression rhythms in 46 human tissues. A stratification by sex and age revealed rich picture of group-specific rhythms, especially in metabolic and cardiovascular tissues that may provide insights into differential disease incidence rates.

## Results

### Comprehensive 24h gene expression rhythms in 46 human tissues

To study the full breadth of rhythmic gene expression programs across the human body, we leveraged Genotype-Tissue Expression (GTEx) bulk RNA-seq measurements, by inferring a circadian reference phase for each donor, called the donor internal phase (DIP). A key property of our approach is that it exploits additional structure in GTEx; one donor provides several tissue samples (typically 10-20) whose phases are therefore not independent. Exploiting this leads to a strongly reduced number of parameters and more robust phase assignments, allowing to cover a significantly larger number of tissues than previous studies [20].

While the time of death (TOD) of donors can be obtained from the GTEx phenotypic data, the relationship to the donor DIPs may be distorted for several reasons: the varying individuals’ chronotypes [3], the accuracy of TODs, the positions in a time zone and ischemic times. In fact, with TODs reference clock genes such as PER2 and NR1D1 exhibited arrhythmic profiles in most tissues (Fig. S1A), and rhythmicity genome and tissue-wide was nearly absent (Fig. S1B). However, clock reference genes (CRG, Methods) showed pairwise correlation structures indicative of a functional clock [21] (Fig. S1C). Thus, TODs are not generally accurately reflecting the internal phases of the donors. Therefore, a comprehensive circadian analysis of GTEx with a high tissue coverage, as well as stratification according to sex and age, requires reliable estimates of circadian phases.

For this purpose, we developed an algorithm to assign DIPs to 914 individuals (Fig. 1A). Briefly, only tissues with more than 48 samples of sufficient RNA integrity and sequencing quality were analyzed (Methods). As we noticed that covariates such as ischemic time, age and sex as well as the type of death (see methods) explained a significant portion of the sample variance, we regressed out those covariates with a linear model and used the residuals for the rest of our analysis (Fig. S1D). To obtain DIPs we proceeded in two steps: first, for each tissue independently, we estimated a tissue internal phase (TIP, referenced to the peak of *PER3* in that tissue) for each sample with CHIRAL. CHIRAL is based on a multivariate harmonic regression that considers twelve CRGs, and in which both the sample phases and genes parameters are fit simultaneously using an expectation-maximization (EM) algorithm (Methods). We validated CHIRAL using time-labeled human samples from muscle [22] (Fig. S1E-F). Secondly, the DIPs were assigned based on the observation that for the majority of donors, the distribution of TIPs across tissues showed a dominant mode, which we robustly estimated and assigned as the DIP for that donor (Fig. S1G-H, Methods). The key advantage of DIPs is that time information is transferred from samples in tissues with a clear temporal structure (high sampling, low confounding sources of variation) to tissues with noisy time signatures. Moreover, DIPs drastically reduce the number of parameters (914 DIPs vs. 16k TIPs). Once DIPs are attributed, we characterized genome-wide rhythmic gene expression programs using standard harmonic regression [23]. Since only one single global reference is needed (we fixed the *PER3* peak expression in muscle to 9am, [22], gene phases can be compared across tissues.

**Figure 1.**
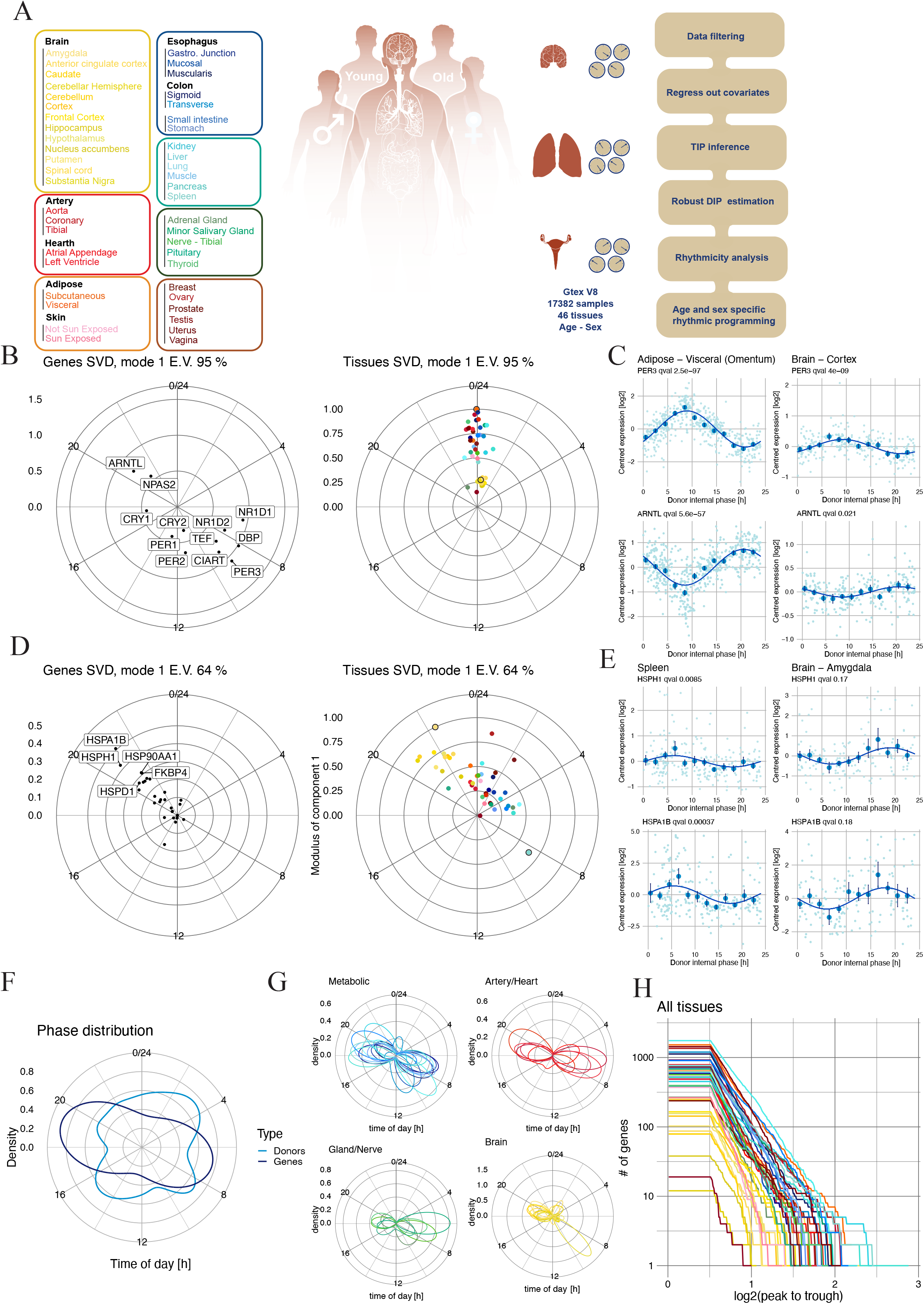
Global circadian ordering of GTEx identifies synchronous circadian clocks in 46 human tissues, as well as extensive biphasic 24h gene expression rhythms. A) Summary of the steps taken to assign one circadian phase (DIP) to 914 donors in the GTEx v8 RNA-seq dataset. This allows us to analyze 46 tissues, and to compare phases of gene rhythms across tissues. Left: Color scheme and list of all tissues present in our analysis. B) First gene (left) and tissue (right) complex vectors from the cSVD performed on clock reference genes (CRGs) show coherent phase and amplitude relationships across tissues (first module captures 95% of the 24h variance, E.V.). The core clock is also tightly synchronized across the human body (spread of ∼2 hours, tissue vector). Brain tissues have the lowest clock gene amplitudes and the adrenal gland peaks earliest. Tissues are color-coded as depicted in A). Polar plots: time runs clockwise, phases have been converted to 24h time. C) mRNA expression levels (log_2_, centered) of two clock genes (*PER3* and *BMAL1*) in two representative tissues. Mean and standard error (SE) computed in 2h bins, and harmonic regression fits are shown (dark blue). D) Heat shock gene module displayed as in B). The genes representing the top 20 HSF1 targets (ChIP-Atlas) show time of day specific expression in all tissues, with higher amplitudes and earlier peak times in brain tissues (yellow). E) mRNA expression levels (log_2_, centered) for two heat shock response genes, represented as in C). F) Polar density plot of the predicted DIPs of the 914 donors and of the peak times of all identified rhythmic genes. G) Phase densities of rhythmic genes (q(BH)<0.2 & log_2_ peak-trough >0.5) for various groups of tissues show biphasic programs, with small rotations. H) Number of rhythmic genes (q(BH)<0.2 & log_2_ peak-trough >0.5) with peak-trough amplitude higher than a threshold (x-axis, log_2_) plotted as a function of the threshold across 46 tissues show different intensities of rhythmic gene expression programs across the human body.

First, we found that DIPs are distributed fairly uniformly along the 24 hour cycle (Table ST1), as the TODs. DIPs and TODs correlated well for fast death donors only (Fig. S1I). However, using the DIPs, *PER2* and *NR1D1* showed robust circadian oscillations using all donors (Fig. S1J, compare with Fig. S1A), indicating that the DIPs capture circadian variability better than TODs. The analysis of CRGs across all tissues showed that while the amplitudes vary, transcript peak times are aligned, the tightest being *TEF* and *ARNTL*, while the most variable is *REVERBA* (*NR1D1*), a gene known to be sensitive to metabolic states (Fig. S1K) [24]. We resorted to the complex-valued singular value decomposition (cSVD), a powerful method to summarize the multi-gene structure of clocks across multiple conditions [25]. The first mode, which captured >95% of the variance, shows that the emerging structure of the human core clock, comprised of two anti-phasic gene groups containing respectively *PER3, DBP*, and *ARNTL, NPAS2* and some genes outside this line such as *NR1D1* and *PER1*, shares similarity with that of other mammals [26,27], and is highly conserved across the 46 tissues (Fig. 1B). Moreover, these clocks are overall well synchronized showing relative offsets of only a few hours. The adrenal gland had the earliest phase, suggestive of the distinct role of adrenal glucocorticoids in systemic clock organization in rats [28]. Tissues showed clear differences in amplitudes, with metabolic tissues (adipose tissue, esophagus, cardiovascular tissues) harboring highest and brain tissues and testis lowest amplitudes (Fig. 1B) [29]. The two highest amplitude clock genes were antiphasic *PER3* and *BMAL1*, showing amplitudes above 8-fold in metabolic tissues, while the oscillations are lower amplitude (<2 fold) but significant in brain tissues (Fig. 1C).

To test whether the DIPs also unveil mRNA rhythms associated with systemic signals, we considered the heat shock stress response genes, known to exhibit rhythmic activity in mice [30]. Heat shock genes (HSF1 targets from Chip-Atlas [31], Methods) showed clear diurnal expression patterns with highest oscillatory amplitude in brain tissues, peaking between 8-10pm near the time of highest body temperature in humans [32] (Fig. 1D). Compared to the clock, we observed a larger spread in peak phases across tissues; *HSPH1* peak times in spleen and amygdala were almost antiphasic (Fig 1E). The high amplitude heat shock program in the brain may reflect a pressure for high proteome integrity in non-renewing tissues [33].

DIPs allow assessing genome-wide rhythmicity across tissues (Fig. S1L, harmonic regression with 24h period on all samples, stratified by Benjamini-Hocheberg adjusted q(BH) statistics, Table ST2). These 24h rhythmic programs showed morning (centered on 7am) and evening (7pm) waves of gene expression throughout the body (Fig. 1F-G), with metabolic tissues harboring the most and brain tissues the least rhythmicity (Fig. 1H). Depending on the tissue, we found between tens and several hundreds of rhythmic transcripts with peak-to-trough amplitudes higher than 2-fold (Fig. 1H). There were slight temporal shifts of those distributions following those of the core clock, with some of the glands being phased earliest, followed by cardiovascular, metabolic and brain tissues (Fig. 1G). Besides CRGs, we found more than 100 transcripts rhythmic in at least 20 tissues, with well-known clock-related genes such as *NFIL3* and *PDK4* as well as glucocorticoid responsive genes such as *FKBP5* and pro-inflammatory cytokine receptors (*IL1RL1, IL1R2*) (Table ST3). 12h ultradian gene expression rhythms, known to exist in mice [34,35], were detected for about 100 genes per tissue (q(BH)<0.2 & amplitude>0.5 log_2_); notable cases were the ovary and the liver whose 12 hour programs were richer (Fig. S1M).

To characterize putative regulatory mechanisms of all rhythmic genes identified, we used the cSVD to analyze experimentally based sets of transcription factor (TF) targets (ChIP-Atlas). We identified putative regulators during the morning and evening waves, involved in immunity, core clock, carbohydrate metabolism and cell proliferation (Fig. S1N). Overall the TFs with the highest explanatory power were the core clock factors CLOCK/BMAL1 (peak target accumulation at 10am), followed by the glucocorticoid receptor (GR) NR3C1 (5pm) corresponding to GR repressed genes [36,37]. In the evening, MYC/MYCN (7pm, cell proliferation), XBP1 (8pm, response to Unfolded Protein Response), and PPARGC1 (8pm, energy metabolism) were activated. Finally, during the night, IRF2 (2am, interferon regulatory factor) and STAT2 (3am, cytokine response) showed their peak of activity (Fig. S1N).

We performed a similar analysis for genes in functional categories (Methods, Table ST4), and first focused on functions showing coherence across many tissues (Fig. S2A). Starting at midnight, we found that immune response genes peak early during the night consistent with the above IRF2 and STAT2 immunity TFs, followed by a response to cholesterol in the early day, coinciding with reported cholesterol peak levels [38]. Around 9am, we observed a peak for caffeine response, and shortly after energy homeostasis, gluconeogenesis, and lipid metabolism genes peaked. Beginning in the early afternoon and extending into the evening, we observed functions related to protein synthesis: amino acid and glucose metabolism, translation, and protein folding. Lastly, cell-cycle related pathways peaked in the evening to late night, coinciding with the predicted MYC/MYCN activities Therefore, pan-rhythmic genes functions in humans consist largely in timed metabolic processes reflecting a switch between low and high energy states during the rest-activity cycle such as lipid oxidation occurring in the morning and synthesis in evening, similar to what is observed during the feeding-fasting cycle in mouse liver (Table ST4) [39]. Among functions that showed higher tissue specificity (Methods), lipid metabolic functions were particularly rhythmic in the liver, amino acid metabolism in the intestine and heat shock response in the brain (Fig. S2B).

### Human sexual dimorphism in circadian rhythms

While human sex-dimorphic gene expression has been studied in GTEx [9], the temporal dimension has not been analyzed. Here, we leveraged the DIP labelling to study sex-dimorphic rhythmic gene expression. First, we compared the core clock in males and females with the cSVD, which showed that the structure of the core clock genes (relative phases and amplitudes) is indistinguishable in the two sexes (the first component captures 93%, Fig. 2A), matching that obtained in the global analysis (Fig. 1B). The phasing of the clocks across tissues remains tight in both male and female, and the clock amplitudes in each tissue were very close (Fig. 2A). While the core clock is invariant and the distributions of DIPs were very similar for males and females (Fig. 2B), clock output programs might differ, which could be masked in the pooled analysis (Fig. 1).

**Figure 2.**
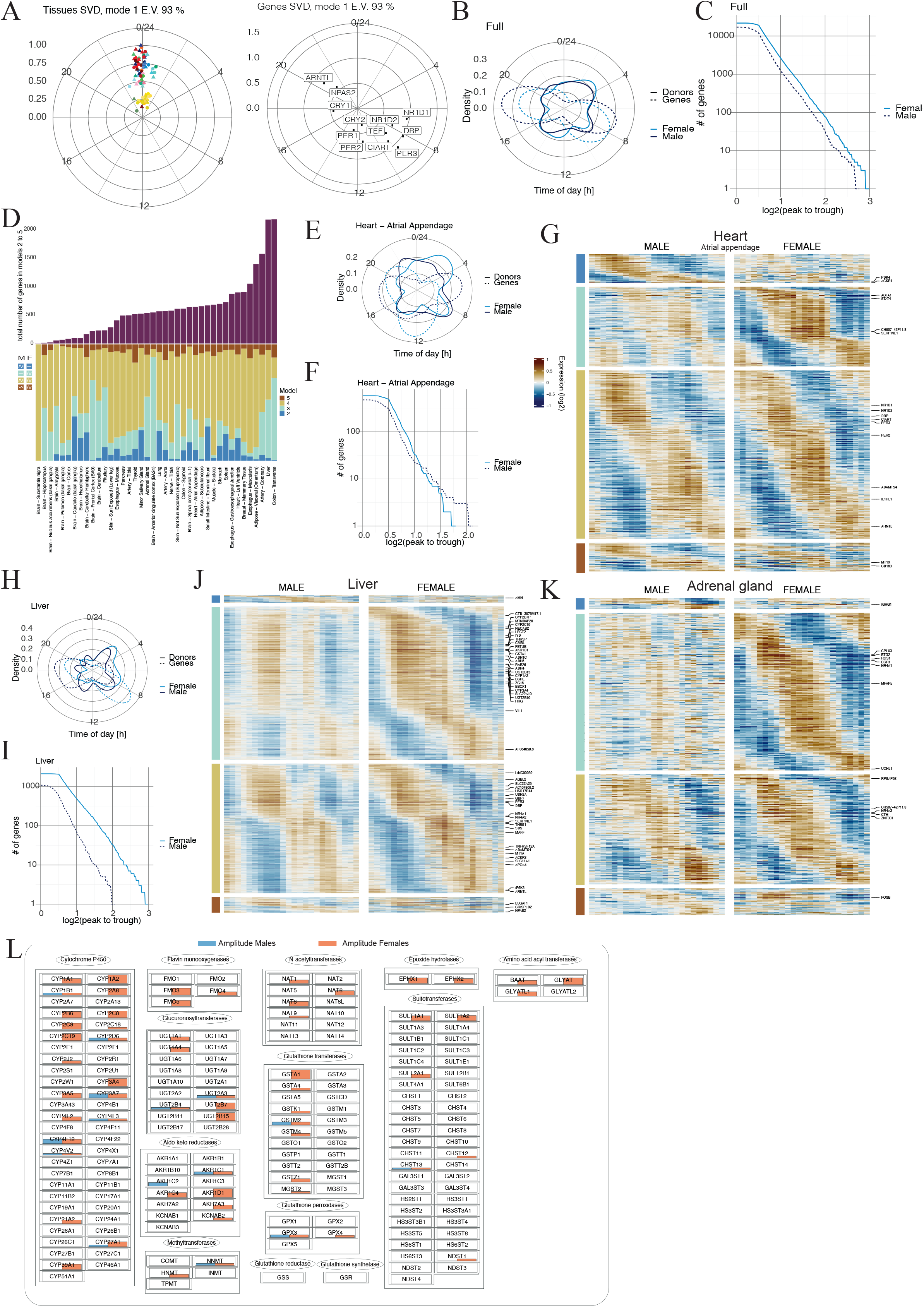
Human sexual dimorphism in circadian mRNA rhythms. A) First gene (left) and tissue (right) vectors of cSVD performed on CRGs indicate invariant clocks between males (triangle) and females (circle). Males and female samples for one tissue were treated as two tissues. The first module captures 93% of the 24h variance, E.V.. B) Polar densities of DIPs (solid line) and gene peak phases (dashed line) in male (dark blue) and female (light blue) donors show morning and evening waves of gene expression. C) Number of 24h-rhythmic genes with amplitude higher than a threshold plotted as a function of the threshold in all tissues combined show a globally higher rhythmicity in females (solid line) than males (dashed line). D) Summary of total number of rhythmic genes in each tissue (top, bordeau) divided according to four statistical models (bottom) show tissue-specificity of sex dimorphic mRNA rhyhtms. Models: 2/blue, 3/cyan, 4/mustard, 5/brick. E) Heart atrial appendage: polar densities of DIPs (solid line) and peak phase of rhythmic genes (dashed line) in for males (dark blue) and females (light blue) show similar overall rhythmicity. F) Heart atrial appendage: number of genes with amplitude higher than a threshold plotted as a function of the threshold for males (dark blue) and females (light blue) reveals an overall similar amount of rhythmic genes in the two sexes. G) Heart atrial appendage: heatmap of mRNA levels for genes in models 2 to 5 illustrates model selection, and shows dimorphic (blue, cyan) and conserved (mustard) rhythmic gene expression. Log_2_ mean centered expression, low/blue to high/brown, 1h bins plotted with a 4h window moving average. mRNAs with top 3% amplitude are explicitly labeled. H) Liver summarized as in E). I) Liver as in F). J) Similar to G). Liver: heatmap of genes in models 2 to 5 recapitulates increased rhythmicity of gene expression in females especially in xenobiotic, fatty acid metabolism and cholesterol related pathways (Fig. S3A). K) Similar to G). Adrenal gland: heatmap of genes in models 2 to 5 shows increased mRNA rhythmicity in females. L) Liver: visualization of biotransformation meta pathway (WP702) colored according to amplitude of genes in males (blue) and females (red) in liver exemplifies the strong female biased rhythmicity in both phase I and phase II enzymes.

To analyze sex-dimorphic gene expression more systematically, we employed a model selection approach to compare rhythmicity across multiple conditions [23,40]. Here we used it to classify each transcript according to five statistical scenarios depending on the rhythmic behavior in both sexes (Methods). The number of rhythmic genes is globally higher in females by a factor of about 30% at all amplitude thresholds (Fig. 2C). Tissues vary in the total number of genes assigned to rhythmic models (Fig. 2D), following the global analysis (Fig. 1H). However, the proportions of genes in the different models unveiled several highly dimorphic tissues such as the adrenal gland and liver showing an excess of female rhythms (Fig. 2D). On the other hand, tissues such as esophagus, skeletal muscle and adipose tissue showed nearly identical rhythms in male and female (Fig. 2D).

We illustrate the model selection with the heart (atrial appendage); while cardiovascular tissues are notable sites of circadian regulation [41] and were previously shown to exhibit circadian rhythmicity in GTEx [20], the stratified analysis reveals a much richer organization. First, we noted that donor phases in the atrial appendage are similar in both sexes and we found no marked difference in the overall number of rhythmic genes or their peak phase (Fig. 2E, F). While the largest proportion of rhythmic genes were shared between male and female, we observed significant rhythmicity that is specific to either sex, and also a group of genes that changes its rhythmic pattern (Fig. 2G).

To illustrate a few specific differences, we found genes related to glucose and fatty acid metabolism (e.g. *PDK4*) to be rhythmic only in males (Fig. 2G, model 2, blue). Clock genes showed similar phase and amplitude across sexes as expected (model 4, mustard). Genes related to metalloproteins (*CD163, MT1X*) and involved in heart disease incidence [42] had different phases or amplitudes between males and females (model 5, brick).

Liver is known for its strong daily rhythms, and marked sex dimorphic gene expression in mammals including human [9]. While dimorphic mRNA rhythms were studied in mice [43], little is known in humans. We found a strong enrichment of mRNA rhythms in females at all amplitudes, mostly as an extensive morning wave of mRNA rhythms (Fig 2H-I). Similar to mice, this enhanced rhythmicity concerned fatty acid and xenobiotics metabolism (Fig. S3A). Moreover, the HSF1 and PPARG transcription factors were predicted to drive female specific rhythms in the liver. These transcription factors are related to heat shock/protein folding response and adipogenesis, respectively and show circadian activation in mice [44], Three pathways stood out as hubs of female specific rhythmicity: the xenobiotic detoxification pathway (WP702), fatty acid breakdown and cholesterol synthesis (Fig S3A). In detoxification, most groups of phase I and II enzymes were strongly enriched in female-specific cyclers (Fig 2L) [45]. In cholesterol biosynthesis, nearly all enzymes, including the rate limiting and statin target *HMGCR* [46], showed marked rhythmicity in females, while these rhythms were strongly damped or absent in males (Fig S3C).

The adrenal gland also exhibited marked female-biased dimorphic rhythmic mRNA levels centered at midday (Fig. 2K). Among those, GR targets were enriched, suggestive of autocrine signaling (Fig. S3B). Moreover, a late evening/early night heat shock stress response is supported both by mRNAs and predicted HSF1 activity. Thus, as glucocorticoid signaling is a systemic synchronizer and organizer of peripheral rhythms [28], the stronger mRNA rhythms in female adrenal gland might corroborate with the overall increased rhythmicity in females at transcriptional and physiological levels [11].

### Age-dependent circadian reprogramming of human gene expression

Next, we analyzed how aging reprograms daily rhythmic gene expression across the human body. Donors were divided in two age groups: younger donors had an age less than fifty, while older donors were above sixty. This division provides a clear separation between age groups and proxy to distinguish pre- and post-menopausal women [47]. We followed the scheme adopted for the sex-dimorphic analysis and showed that the overall amplitudes, phases and relative relationships of the CRGs were conserved with age (Fig. 3A). Indeed, intra-tissue differences between the older and younger samples were negligible for the CRGs (Fig. 3A). In contrast, rhythmic programs overall were strongly dampened in aged donors, with almost a 10-fold decrease in the number of rhythmic genes with amplitudes higher than two log_2_ units (Fig. 3B). Moreover the existence of two waves of gene expression is conserved across age groups (Fig. 3C). Using the model selection approach, we noticed that this trend of loss of rhythmicity with aging occurs in the vast majority of tissues throughout the body (Fig. 3D).

**Figure 3.**
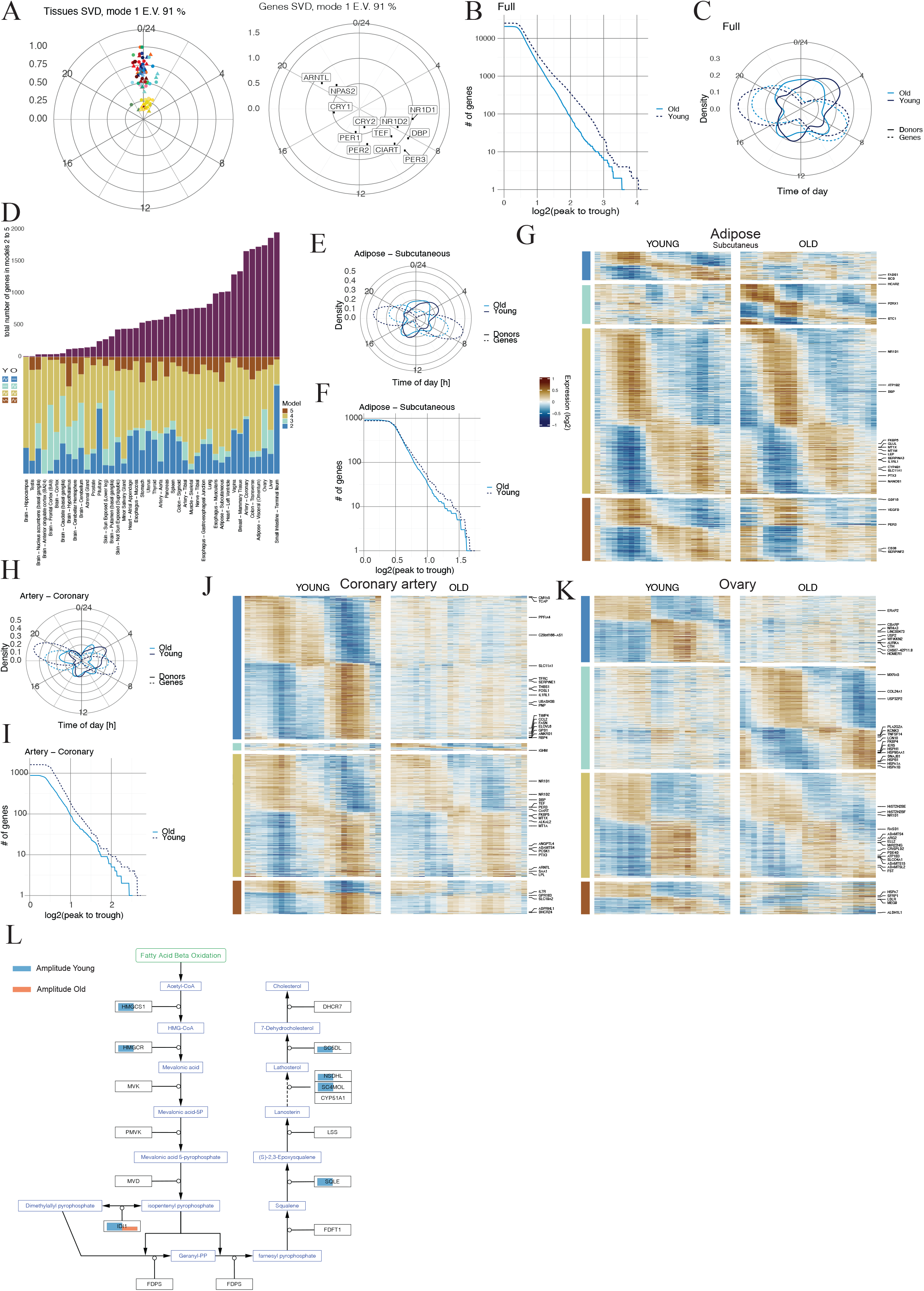
Age-dependent circadian reprogramming of human gene expression. A) First gene (left) and tissue (right) vectors of cSVD performed on CRGs indicate invariant clocks between younger (triangle) and older (circle) donors. Males and female samples for one tissue were treated as two tissues. The first module captures 91% of the 24h variance, E.V.. B) Number of 24h-rhythmic genes with amplitude higher than a threshold plotted as a function of the threshold in all tissues combined exhibit substantially stronger 24h mRNA rhythmicity in younger (dark blue) compared to older (light blue), especially for higher amplitudes. C) Polar densities of the DIPs (solid line) and gene phase (dashed line) for younger (dark blue) and older (light blue) donors. D) Summary of the number of rhythmic genes in each tissue divided according to the model selection approach recapitulates the global behavior of tissues (metabolic tissues are more rhythmic, brain less) and the general trend of loss of rhythmicity as a result of aging. Models: 2/blue, 3/cyan, 4/mustard, 5/brick. E) Polar densities of DIPs (solid line) and peak phase of rhythmic genes (dashed line) in adipose subcutaneous for younger (dark blue) and older (light blue) show a decay in the biphasic rhythmic programming as age progresses. F) Adipose subcutaneous: number of genes with amplitude higher than a threshold plotted as a function of the threshold for young (dark blue) and old (light blue) shows highly conserved rhythmic programs with only a mild dampening of high amplitude rhythms in elderly. G) Adipose subcutaneous: Heatmap of genes in models 2 to 5 highlights the mainly conserved rhythmic programming (mustard) as well fewer losses (blue) and gains (cyan) with aging. Log2 mean centered expression, low/blue to high/brown, 1h bins plotted with a 4h window moving average. mRNAs with top 3% amplitude are explicitly labeled. H) Coronary artery: polar density plot of DIPs (solid line) and peak phase of rhythmic genes (dashed line) for young (dark blue) and old (light blue) shows morning and evening waves of gene expression that decay with age. I) Coronary artery: number of genes with amplitude higher than a threshold plotted as a function of the threshold for young (dark blue) and old (light blue) emphasizes more abundant rhythmic programming in younger donors across all amplitudes. J) Similar to G). Coronary arteries: heatmap of genes in models 2 to 5 illustrates strong morning and evening waves of gene expression and exemplifies instances of loss of functional rhythmic programs. K) Similar to G). Ovary: heatmap of genes in models 2 to 5 depicts differential rhythmic programming of gene expression as a function of age. L) Artery coronary: visualization of cholesterol biosynthesis pathway (WP197) colored according to amplitude of genes in young (blue) and old (red).

However, not all tissues show a decay in rhythmic mRNAs. In fact, adipose tissues, esophagus and skeletal muscle showed highly conserved rhythmicity across age, with a majority of genes exhibiting statistically identical rhythms in the two groups (model 4, mustard, Fig. 3D). This is well illustrated in adipose subcutaneous, in which the morning and evening waves are pronounced in both younger and older donors (Fig. 3E), with a vast majority of invariant gene rhythmicity, as well as fewer gains and losses with age (Fig. 3F, G).

We next focused on coronary arteries, a tissue with strongly age-biased rhythms. While morning and evening waves were observed in both groups (Fig. 3H), the number of rhythmic mRNAs in elderly was only about half that of younger donors across all amplitudes (Fig. 3I). The fraction of transcripts classified as rhythmic only in the younger donors shows that a significant fraction of expression programs lose rhythmicity as age progresses (model 2, blue, Fig. 3J). Important programs that lose rhythmicity include cholesterol biosynthesis, FA synthesis, and the glycolysis regulation (Fig. 3L, S4A), processes known to be deregulated in vascular smooth muscle cells in cardiovascular diseases [48]. Strikingly, most enzymes in the cholesterol biosynthesis pathway, including *HMGCR*, were rhythmically expressed in young coronary arteries but lost this feature with age (Fig. 3L). Other high amplitude genes relevant for cardiovascular pathophysiology include the fatty acid elongase *ELOVL6*.

Interestingly, we observed marked differences in ovaries between pre- and post-menopausal women (Fig. 3K), consisting of both lost and gained mRNA rhythms. While rhythmicity in lipid and cholesterol biosynthesis was suppressed in aged donors, as in the coronary arteries (Fig. S4B), stress and in particular heat shock response genes became highly rhythmic, as also supported by the predicted HSF1 transcription factor activity (Fig. S4B-C). Though this signature of a putative thermal stress response in post menopausal women was found only in ovaries, it may reflect circadian patterns of core body temperature in post-menopausal women [49].

Lastly in some tissues, we uncovered a scenario whereby genes that lost 24h rhythmicity switched to a 12h ultradian periodicity in the elderly. This is clearly exemplified by the pituitary gland, the liver and colon where 12h rhythms arise in 30-50% of genes classified as only rhythmic in young (Fig. S4D-F). These tissues are regulators of human physiology driving the rhythms of temperature, energy metabolism and absorption, and indirectly influencing the sleep wake cycle [50]. Such destabilization of 24h periodicity in favor of an ultradian state as a result of aging might reflect differences in reception of Zeitgeber cues in function of age [51].

## Discussion

With the inferred DIPs, we showed that the phase and amplitude relationships of CRGs are well conserved across tissues, age groups and sexes. Moreover, clocks were largely in phase in the 46 tissues analyzed, with the adrenal gland peaking earliest. The concomittent signature of a seizable wave of negative GR targets in the afternoon suggests a key role for released adrenal glucocorticoids for human body-wide circadian gene expression rhythms.

Among other TFs explaining a significant portion of the body’s 24h rhythms, HSF1 target heat shock genes showed differential rhythmic behaviour across a variety of tissues in function of sex and age. Interestingly, HSF1 activity showed high-amplitude oscillations in several elderly tissues (adipose-subcutaneous, adipose visceral, brain frontal cortex, nerve-tibial, stomach), which was particularly noticeable in the ovary. There, the oscillatory patterns of heat shock response genes replace those of lipid and cholesterol metabolism genes which are flat in aged donors, possibly as a result of menopause.

The striking observation that rhythmic liver gene expression was significantly more prevalent in females may bring new insights into non-alcoholic fatty liver disease, where incidence is almost double in males [52]. While it would need to be established that mRNA rhythms propagate to rhythmic enzymatic activities, we may speculate that enhanced rhythmicity in metabolic pathways may be an attenuating risk factor for certain diseases. Such strong differential gene expression rhythmicity, in particular in xenobiotic detoxification genes, may also provide opportunities to develop sex-specific chronopharmacology [53]. Similarly, as cardiovascular diseases show known age-dependent incidence rates, in fact enough to characterize age as a direct risk factor [54], the loss of nearly 800 mRNA rhythms in coronary arteries in the older age group could explain the higher prevalence in elderly. The strong difference in rhythmic processes could also be relevant to explain the uneven timing of sudden cardiac death across sexes and ages and possibly open new avenues for prevention [55] and effectiveness of timed medication [56]. Lastly SERPINE1, a gene needed for controlled blood clot degradation [57,58], was arrhythmic in elderly or in male cardiovascular tissues; thus strategies based on restoring such rhythms may help prevent cardiovascular diseases showing lower incidence in young and in females.

In conclusion, the possibility to generate sufficient comprehensive and suitably stratified views of temporal gene expression rhythms accross the human body may help the optimal timing and dosing of drugs, notably according to sex and age, both for improving the efficacy or reduce side effects [59].

## Acknowledgments

The Genotype-Tissue Expression (GTEx) Project was supported by the Common Fund of the Office of the Director of the National Institutes of Health, and by NCI, NHGRI, NHLBI, NIDA, NIMH, and NINDS. The data used for the analyses described in this manuscript were obtained from the GTEx Portal on 04/12/20 and dbGaP accession number phs000424.GTEx.v8.p2.c1.GRU on 03/30/2022. This project was funded by a Swiss National Science Foundation individual project grant 310030B_201267 to F.N. We thank Frédéric Gachon for insightful discussions.

## Methods

### RNA-seq data preprocessing

Gene read count data from GTEx V8 (2017-06-05_v8_RNASeQCv1.1.9) were downloaded. For each tissue, genes with less than 10 counts on average were discarded. Counts were normalized by library size and scaled using the TMM method from edgeR [60], and then converted to counts per million (CPM). Normalized counts were then log-transformed (log_2_) after addition of a pseudo-count of 2. Genes with on average less than 1 CPM per issue were discarded. To control for sample quality in the first step of the analysis, samples with RIN numbers larger than 0.6, total number of reads larger than 40 millions, mapping quality larger than 80%, a positive ischemic time and known autolysis score were selected. Tissues with less than 48 samples were discarded. Overall, those initial filtering steps led to 35 tissues comprising about 10’000 samples (set 1) that were used for the TIP/DIP assignments. Note that once the DIPs are assigned, these criteria will be relaxed to include additional samples and cover in total 16’000 samples across 46 tissues (set 2).

Using principal component analysis PCA for data exploration, we noticed that several covariates available as metadata explained significant portions of the variance (Fig. S1D). Therefore, we regressed out ischemic time, sex, age and type of death as categorical variables with a linear regression model on the normalized count data. The residuals of these fits were then used for all subsequent analyses. We note that by linearly removing sex and age contribution we are just removing the mean value and this does not influence the possible differences in rhythmicity.

### Inference of DIPs

First, we assigned a TIP for each sample in each tissue using CHIRAL (see below and Supplemental Method). This first step is similar to that in [20], except that data are preprocessed which reduces the influence of confounding variance sources. The key conceptual advance is the following: due to the nature of the GTEx the TIPs for one donor are not independent but have additional structure. At the level of TIPs we have a set of internal phases φ_*t,d*_ that depend on the donor (*d*) and tissue (*t*). Our main assumption consists in expressing the TIPs as a sum of two contributions: φ_*t,d*_ = φ_*d*_ + δφ_*t*_, where φ_*d*_ is the donor specific phase (DIP) and δφ_*t*_ is the tissue specific shift which is assumed to be common to all donors in a set of samples (as a convention we use skeletal-muscle as the reference, i.e. δφ_*t*=*skeletal muscle*_ = 0). This drastically reduced the number of parameters, from |*d*| × |*t*| to |*d*| + |*t*| which renders the whole procedure more robust. To identify the DIPs from the TIPs we then applied a heuristic based on the following (using the samples in set 1). We observed that the distributions of φ_*t,d*_across tissues for one donor were fairly peaked (Fig. S1G-H), indicating that the majority of tissue harbor very similarly phased clocks. The outliers were generally due to an inherent instability of phase ordering algorithms that is linked to the collinearity of clock transcripts (most transcripts peak in the *BMAL1* or almost exactly opposite *PER3* phase), and which is particularly present in tissues with weak or confounded clocks. We thus defined the robustly estimated mode of this peaked distribution as the DIP. Specifically, for each donor, after identifying the dominant mode we computed the circular mean of samples within a 2h interval. In this approach tissues with weak clocks, or tissues which contain significant sources of non temporal gene expression variability do not bias the donor phase estimates. Moreover, unlike time of death (TODs), DIPs do not need to be corrected for chronotype as they directly measure the internal clock phase.

Finally, after DIPs are assigned for all donors in set 1, we relaxed our sample selection filter by only selecting for RNA quality (RIN > 0.4). This allowed us to expand the analysis to tissues and samples of lower data quality from which a robust circadian phase could not have been inferred alone. Overall, this allowed us to study the daily oscillations in 46 tissues and 16k bulk RNA-seq samples from GTEx (set 2).

### Estimation of TIPs with CHIRAL

As a first step in the algorithm, we reconstruct internal circadian phases for samples in a given tissue (tissue internal phases or TIPs), which we then improve as described above. For this purpose, we model a gene expression matrix (*E*_*gc*_, log transformed gene expression measurements) using a multivariate harmonic regression model *E*_*gc*_ = *μ*_*g*_ + *a*_*g*_ *cos* (φ_*c*_) + *b*_*g*_ *sin* (φ_*c*_) + ε for conditions c and genes g with Gaussian noise ε. If a gene does not carry time information in that tissue, the corresponding gene Fourier parameters (*a*_*g*_, *b*_*g*_) can be very small or even zero (which we consider as two scenarios by introducing a two state mixture model for each gene). In fact we treat the model probabilistically with prior Gaussian distributions over the gene parameters and using a Bayesian calculation to marginalize these and derive a posterior distribution over the unknown phases φ_*c*_. When then derive maximum a posteriori estimators using an expectation maximization (EM) algorithm that has a similar structure as probabilistic PCA [61], but with an additional constraint that each 2D cosine and sine vector has norm 1, which we impose exactly using lagrange multipliers. We set a stopping condition when the maximum absolute phase difference was smaller than 0.001 radians between two subsequent iterations of the EM. The initial phases are seeded using a mean field approximation of a simplified model, which yields a so-called XY spin glass, and which can be solved using a simpler iterative scheme (Supplemental Methods).

### Harmonic regression

One we assigned the DIPs φ_*d*_ to all donors in set2, we assessed rhythmicity and inferred gene coefficients in each tissues using harmonic regression as previously *E*_*gd*_ = *μ*_*g*_+*a*_*g*_ *cos* (φ_*d*_) + *b*_*g*_ *sin* (φ_*d*_) + ε [25]. Rhythmicity was assessed using a likelihood ratio test between a rhythmic and a flat (*a*_*g*_ = *b*_*g*_ = 0) model. P-values were computed from a chi-squared distribution. The complex representation of a gene used in the cSVD technique is *a*_*g*_ + *ib*_*g*_.

### Clock correlation matrix

We computed a clock correlation matrix following the ideas of [21]. A matrix of correlation of the CRGs allowed us to verify if there was a clock in a tissue. We wanted one matrix to represent the ensemble of tissues to understand if a clock is generally present in the GTEx data; so, we took a weighted average of the various tissue specific correlation matrices weighted by the number of samples in that tissue. We found that indeed the structure of the clock is generally present in the GTEx dataset (Fig. S1C).

### Model selection approach

To study the stratified scenarios (male vs. female and age groups), we first selected possible rhythmic genes as those which had a q-value (Benjamini Hochberg, BH) < 0.2 and a log_2_ peak-to-trough amplitude > 0.5, when performing harmonic regression with either all samples, or only samples in of the two conditions. On this set of genes, we adopted a model selection approach (DryR) similarly to (Atger, 2015; Weger, 2021) using the Akaike Information Criterion (AIC), resulting in 5 different models. Model 1 (never shown) represents genes flat in both conditions, male-female or young-old. Model 2 (blue) represents genes only rhythmic in the first condition, male/young, and model 3 (cyan) those in the second condition, female/old. Model 4 (mustard) is composed of genes rhythmic in both conditions with the same parameters (phase, amplitude) while in model 5 (brick), genes are rhythmic in both conditions with different parameters. As the number of samples in the two compared groups might differ, we subsampled the group with more samples to have a model selection without biases due to sample sizes.

### Complex-valued Singular Value Decomposition (cSVD)

We used the complex-valued singular value decomposition (cSVD) to decompose and represent rhythmic gene expression across multiple tissues as in [25]. The algebra of complex numbers and linear mathematics of cSVD is optimally suited to decompose oscillatory gene expression matrices into rank 1 modules consisting of complex valued gene and condition vectors allowing to display the relative phases and amplitudes in genes space, and how these genes modules are scale and phase shifted in tissues (visualized in two separate polar plots). Here we mostly display the rank one approximations and indicate the fraction of the variance explained by those. In the representations, we break intrinsic symmetries by setting the maximum amplitude of the tissue vector amplitude to one, the eigenvalue of the first mode to one, and the mean phase of the tissue vector to zero. This allows the gene representation to correctly show both amplitude and phase of the genes. For each relevant gene set (CRGs, GO gene sets, TF targets, etc), we constructed *N*_*gene*_ × *N*_*tissue*_ matrices containing the Fourier coefficients (*a*_*g*_ + *ib*_*g*_) in complex notation. The first left (gene) and right (tissue) complex vectors of the cSVD were used to calculate the relevant metrics.

### Gene Ontology analysis with cSVD

We performed cSVD with gene sets representing all GO terms or Wikipathways [62] to identify commonly rhythmic (Fig. S2A) and tissue-specific (Fig. S2B) functions. To prevent high variance genes with poor sinusoidal behavior from dominating, for this analysis, we renormalized the complex matrix entries such that the amplitudes reflect the fraction of explained variance by the harmonic fit for each gene in each tissue. This analysis using the cSVD allowed us to calculate a number of useful metrics for each GO term (see below for a detailed list) and used to make selections.

Commonly rhythmic functions in (Fig. S2A) have been selected due to either a high variance explained (variance) by the first SVD component, a high phase coherence of the mean peak time across tissues (tissue_phase_similarity), or a high coherence of the peak times of the genes across tissues (gene_phase_similarity). In particular, we ordered the GO terms according to these three metrics and extracted the top 50 from each. Then we removed redundancies and eliminated general GO terms to reach the list in Fig. S2A. Finally, by inspecting all genes in these functions we coarse grained the terms into functional themes for easier interpretation. These gene functions are displayed as ordered by (mean) time of day. To identify more specific circadian tissue effects, we filtered functions according to either a high entropy of the first tissue vector (tix_entropy), or a high standard deviation of the mean peak phase across tissues (tissue_phase_similarity) (Fig. 2B). As before, we ordered the GO terms according to these three metrics and extracted the top 50 from each. Then we removed redundancies and eliminated general GO terms to remain with the list in Figure S2B. The coarse grained themes often overlap between Figures S2A and S2B, however, the specific pathways or genes involved in these functions are often not the same.

#### Metrics calculated for each GO or Wikipathway gene set

*Variance*: variance explained by the 1st cSVD component.

Mean_gene_time: argument of the sum of complex valued gene parameters of 1st cSVD component.

*Mean_ampl*: mean of absolute values of complex valued gene parameters of 1st svd components.

*Tix_entropy*: entropy of the probability distribution given by the modulus squared of the complex valued tissue parameters of 1st cSVD component

*Gene_phase_similarity*: modulus of the sum of complex valued gene parameters of 1st cSVD component.

*Tissue_phase_similarity*: modulus of the sum of complex valued tissue parameters of 1st cSVD component.

### GO analysis enrichment

For the functional analysis of the stratified analysis (male vs. female and age group), we used the “enrichR” R package providing an interface to the Enrichr database [63]. For each model, we performed enrichment analysis for three databases: GO_Biological_Process_2021, KEGG_2021_Human, WikiPathway_2021_Human. Terms with an adjusted p-values smaller than 0.1 and a combined score larger than 20 were selected. Duplicated terms with a similar list of genes were discarded. The top fifteen terms (i.e. smallest adjusted p-values) across databases are reported in the heatmaps (e.g. Fig. S3A).

### Transcription factor (TF) analysis with cSVD

We performed a similar cSVD analysis as that of GO terms on sets of transcription factor (TF) target genes, which allowed us to visualize the predicted peak phase activities of each TF throughout the day (Fig. S1N). From the human Chip-Atlas [31], we selected for each transcription factor the top 100 targets and performed cSVD on them (supplementary table ST4). This allowed us to calculate the same metrics as for the GO term analysis for each TFs. As for the GO, we used the renormalized Fourier coefficients to account for explained variance.

### TF analysis with MARA

This approach was used for the stratified analyses (male vs. female and age group). ChIP-Seq target gene data were downloaded from ChIP-Atlas for the H. Sapiens (hg38) genome [31]. Peak-calls overlapping a 5kb window around the TSS of a gene were assigned to it. For each sample, MACS2 score was normalized to its maximum value. For each transcription factor (TF), the normalized MACS2 score was averaged across all samples. To infer TF activity, we adapted a penalized regression model [64] as previously [25]. In the stratified analysis, for each statistical model, we inferred TF activity using mean-centered mRNA expression level and ChIP-Seq data described above. We performed model selection on the inferred TF activities and selected TFs classified in the same model with high confidence (AICW > 0.5). The top 10 TFs in terms of amplitude or explained variance (z-score) were shown in the heatmaps.

### Gene lists

*Clock reference genes (CRG)*: *DBP, PER3, TEF, NR1D2, PER1, PER2, NPAS2, ARNTL, NR1D1, CRY1, CRY2, CIART*.

*Heat shock response gene*s (top HSF1 target from the human ChIP-Atlas): *HSPE1, HSPD1, HMGB1, USPL1, HSP90AA1, DNAJB6, XPNPEP3 ST13, HSPH1, MSH3, HSPA8, TCP1, MRPL18, MTRNR2L8, MTRNR2L2, FKBP4, HSP90AB1, MAMDC2, COMMD1, CCT4, UBB, RGPD2, CCDC117, STIP1, HSPA1B, MTRNR2L1, CHORDC1, APTX, DNAJA1*.

## Supplemental Tables

**Table ST1**. Donor internal phases (DIPs) for GTEx samples. SUBJ.ID: unique GTEx donor identifier; Phi: DIP in radians; Hour: DIP*12/pi to report DIPs on the 24h scale.

**Data Tables ST2**. Zip of txt files with names tissue_name.txt. Each file has genes as rownames. The first N column names are the unique GTEx sample identifiers. Values in these first N columns contain our processed mRNA expression levels. The last 38 columns contain statistical variables that we calculated on the 24h gene expression rhythms. These are expressed in the form *variable*.*stratification*.

*Stratifications:*

ALL: all samples fit together

YOUNG/young: only young samples

OLD/old: only old samples

Males/MA: only male samples

Females/FE: only female samples

MF: Male-female comparison

AGE: young-old comparison

*Parameters:*

pval: p-value qval/qvals: q-value(BH)

amp: amplitude peak to trough phase: peak phase (24h scale)

a,b: gene Fourier coefficients of the harmonic regression

model: integer value between 1 and 5, corresponding to the 5 models of our model selection approach

AICW: Weight of the top model following Akaike Information Criterion relamp: relative amplitude

**Table ST3**. Gene and tissue rhythmicity. Binary matrix indicating whether mRNAs are rhythmic in the tissues. Criterion is that used in the selection for Figure 1 (q(BH)<0.2, amplitude >0.5 log2).

**Table ST4**: Gene Ontology and Transcription Factor analysis with cSVD.

Extra var: excess variance with respect to a relationship variance explained vs. log(#genes).

*Variance*: Fraction of the variance explained by the 1st cSVD component.

*Mean_gene_time*: argument (angle) of the sum of complex valued gene parameters of 1st cSVD component.

*Mean_ampl*: mean of absolute values of complex valued gene parameters of 1st SVD components.

*Top_eigenvalue*: maximum eigenvalue of the cSVD decomposition.

*Scaled_EV*: top eigenvalue divided number of genes.

*N_genes*: the number of genes.

*Other columns*: tissue names. Entries are the absolute value of the complex tissue vector scaled such that the maximum is 1.

**Figure S1.**
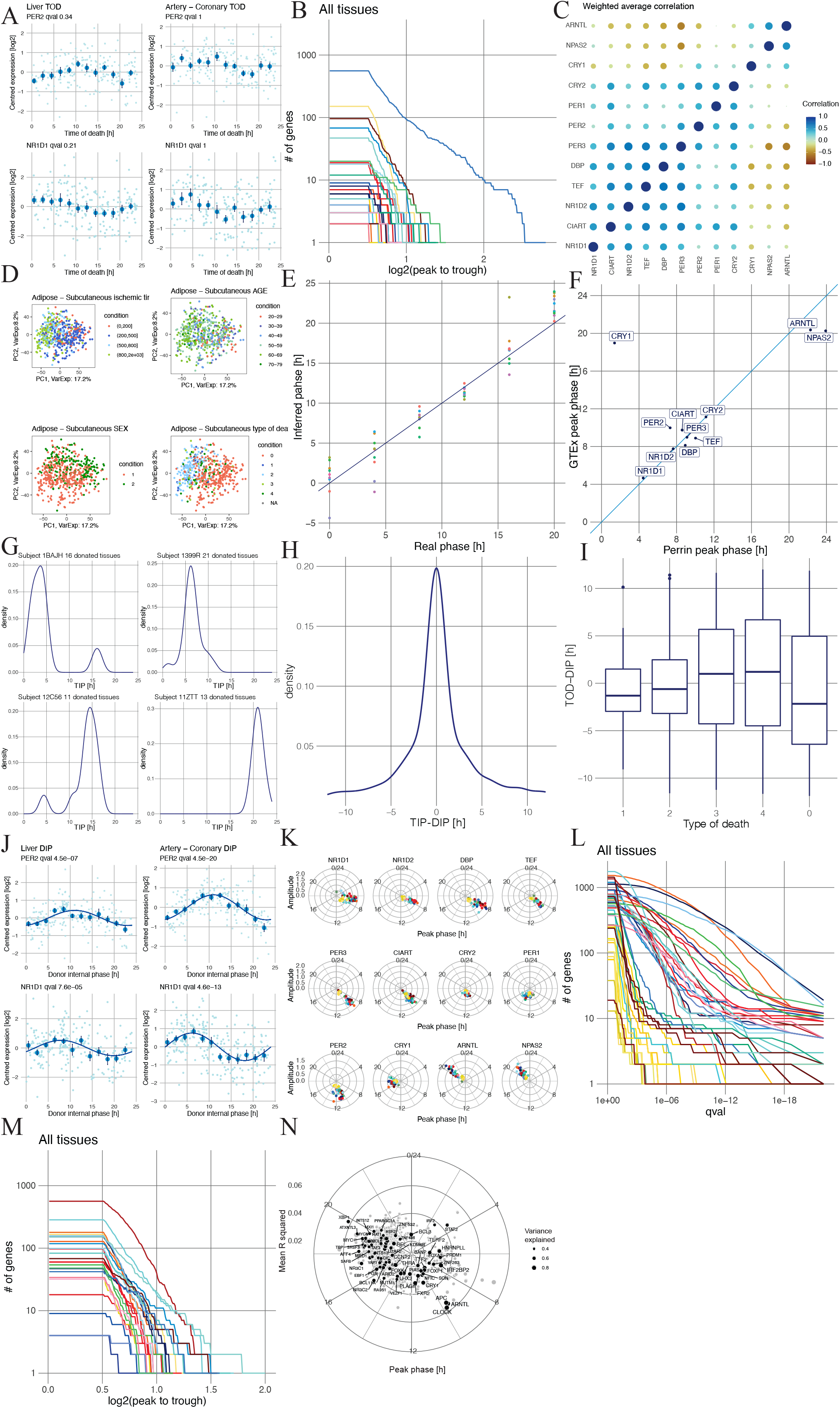
Characteristics and validation of phase inference method for DIPs. A) Clock genes PER2 and NR1D1 exhibit statistically arrhythmic profiles in metabolic tissues as a function of GTEx time of death (TOD). Log_2_ centered expression for each donor (light blue). Mean and standard error (SE) computed in 2h bins are also shown (dark blue). B) Number of rhythmic genes (q(BH)>0.2 & peak-trough difference >0.5, same criteria as in Fig. 1) with peak-trough amplitude higher than a threshold (x-axis, log_2_) plotted as a function of the threshold across 46 tissues shows very low rhythmicity across the body using indicated time of death. C) Weighted correlation matrix (Methods) of CRGs with two correlated blocks of genes supports the integrity of the clock across the body. D) Principal Component Analysis (PCA) analysis of samples for adipose subcutaneous in PC1-PC2 space color coded according to different meta variables calls for removal of undesired sources of variability in the data. PC1 and PC2 explain respectively 17.2% and 8.6% of the total variance. E) Phase inferred using CHIRAL on time labeled human muscle data [22] shows good performance with a median absolute error of 1.02h. F) Comparison of peak phases of CRGs in labeled data [22] and GTEx muscles. G) Illustrative TIP distributions with one clear peak suggest the possibility of inferring the DIP. H) Distribution of the TIP (shifted by the DIP) for all samples and donors shows a clear and tight peak. I) Boxplot of the difference between TOD and DIP as a function of type of death (hardy scale) shows higher concordance for fast (types 1, 2) compared to slow (types 3, 4, 0) deaths. J) Clock reference genes PER2 and NR1D1 exhibit robust oscillatory behavior in metabolic tissues as a function of DIP. Mean and standard error (SE) computed in 2h bins, and harmonic regression fits, are also shown as in A). K) With the DIP, clock gene phases can be compared across tissues. Phases (clockwise) and amplitudes (radial) of CRGs across tissues highlight overall tight phases. NR1D1 is the most variable gene. Tissues are color coded as in Fig. 1A. L) Number of rhythmic genes (q(BH)>0.2 & log_2_ peak-trough difference >0.5) with q-value smaller than a threshold (x-axis) plotted as a function of the threshold across 46 tissues shows numerous and highly significant rhythms in the metabolic tissues. Tissues are color coded as in Fig. 1A M) Number of 12h-rhythmic genes (q(BH)<0.2 & log_2_ peak-trough difference >0.5) with peak-trough amplitude higher than a threshold (x-axis, log_2_) plotted as a function of the threshold across 46 tissues show different intensities of rhythmic gene expression programs across the human body. N) Transcription factor activities. Polar plot of mean phase and mean R^2 values for the top 100 targets of all transcription factors (ChIP-Atlas) across the body obtained via cSVD (Methods, Table ST3). These factors are mostly one with pan-rhythmic activity, i.e. the target genes are rhythmic in multiple tissues. Note: The APC antibody used in those experiments (Bethyl Laboratories # A300-981A) significantly crossreacts with CLOCK.

**Figure S2.**
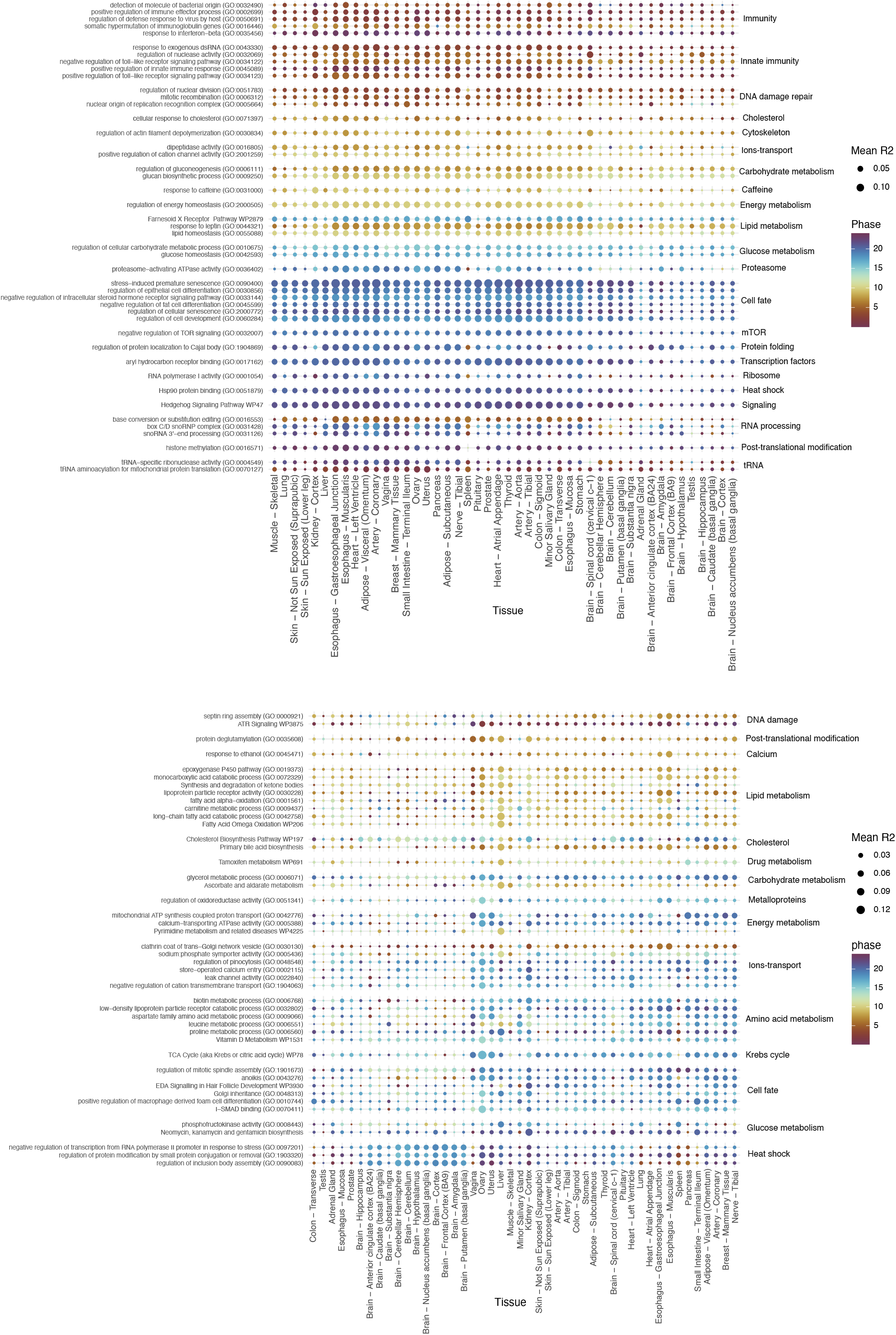
Pan rhythmic and tissue biased GO terms and Wikipathways. A) Set of GO terms showing a coherent rhythmicity across the body in the cSVD approach (Method). Size of the circle is the mean R squared (harmonic regression) of the genes, color is the mean phase. B) Set of tissue specific rhythmic GO terms highlight enhanced rhythmicity of some function in specific tissues. Representation as in (A).

**Figure S3.**
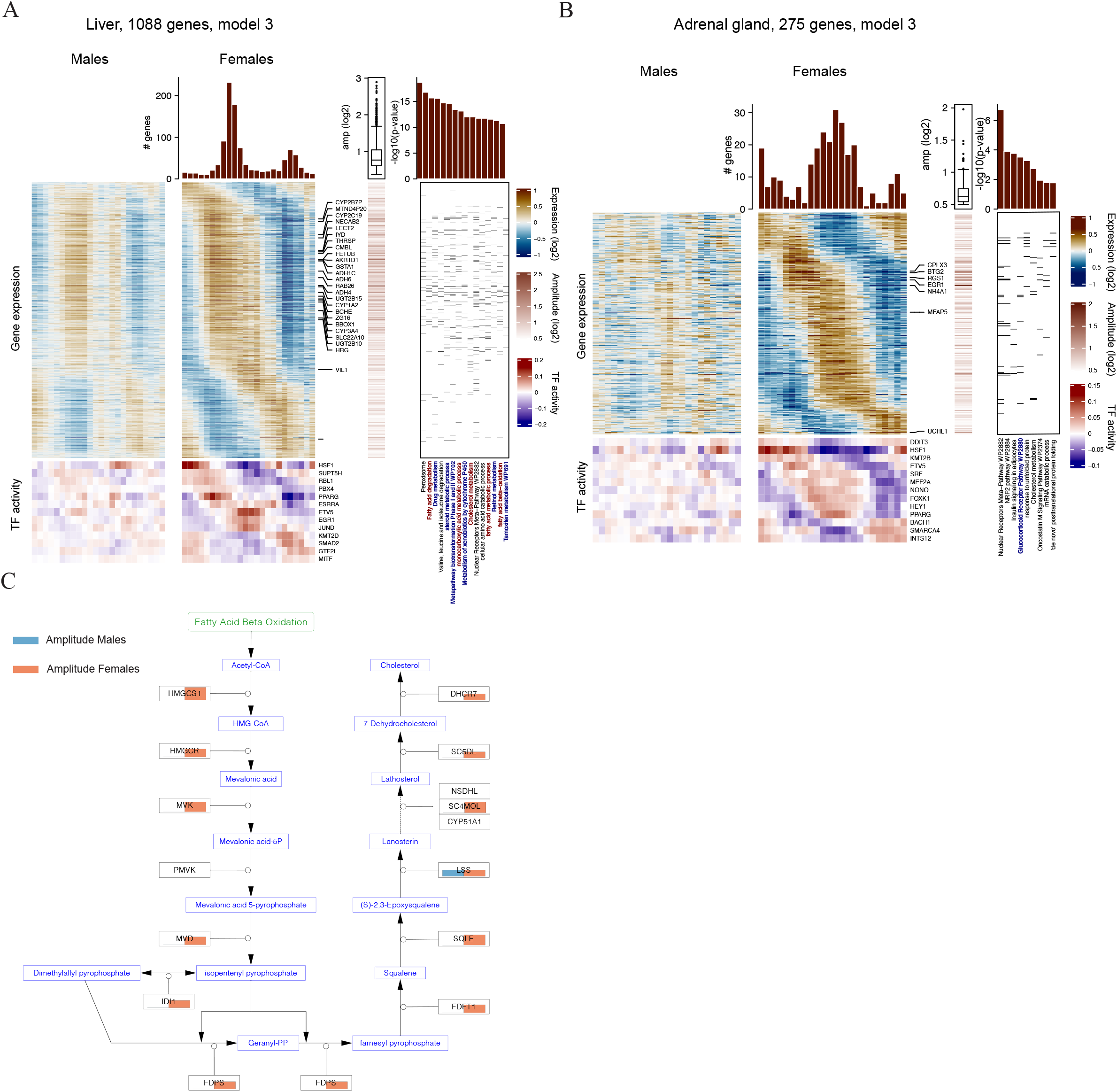
Compound heatmap visualizations of sex dimorphic rhyhtmic mRNA programs in selected tissues. A) Female-biased liver mRNA rhythms. Left part: Heatmaps (model 3) for genes being more strongly rhythmic in female liver (left heatmap: males; right heatmap: females, blue/brown scale). mRNAs with the top 3% amplitudes are explicitly labeled. The heatmaps show log_2_ mean centered expression, low/blue to high/brown, 1h bins plotted with a 4h window moving average. Corresponding transcription factor activities (MARA, Methods) are shown below (blue/red scale). Right part: GO and WikiPathway enrichments. P-values, TF activities, amplitudes of gene fits are also reported (Methods). Fatty acid and cholesterol metabolism are written in red and blue, respectively. B) Genes only rhythmic in the female adrenal gland displayed as in A). Note the enrichment of GR signaling genes (blue text). C) Cholesterol biosynthesis pathway (WP197) colored according to the oscillatory amplitudes of genes in males (blue) and females (red) in the liver, shows that all enzymes but one are rhythmic in females.

**Figure S4.**
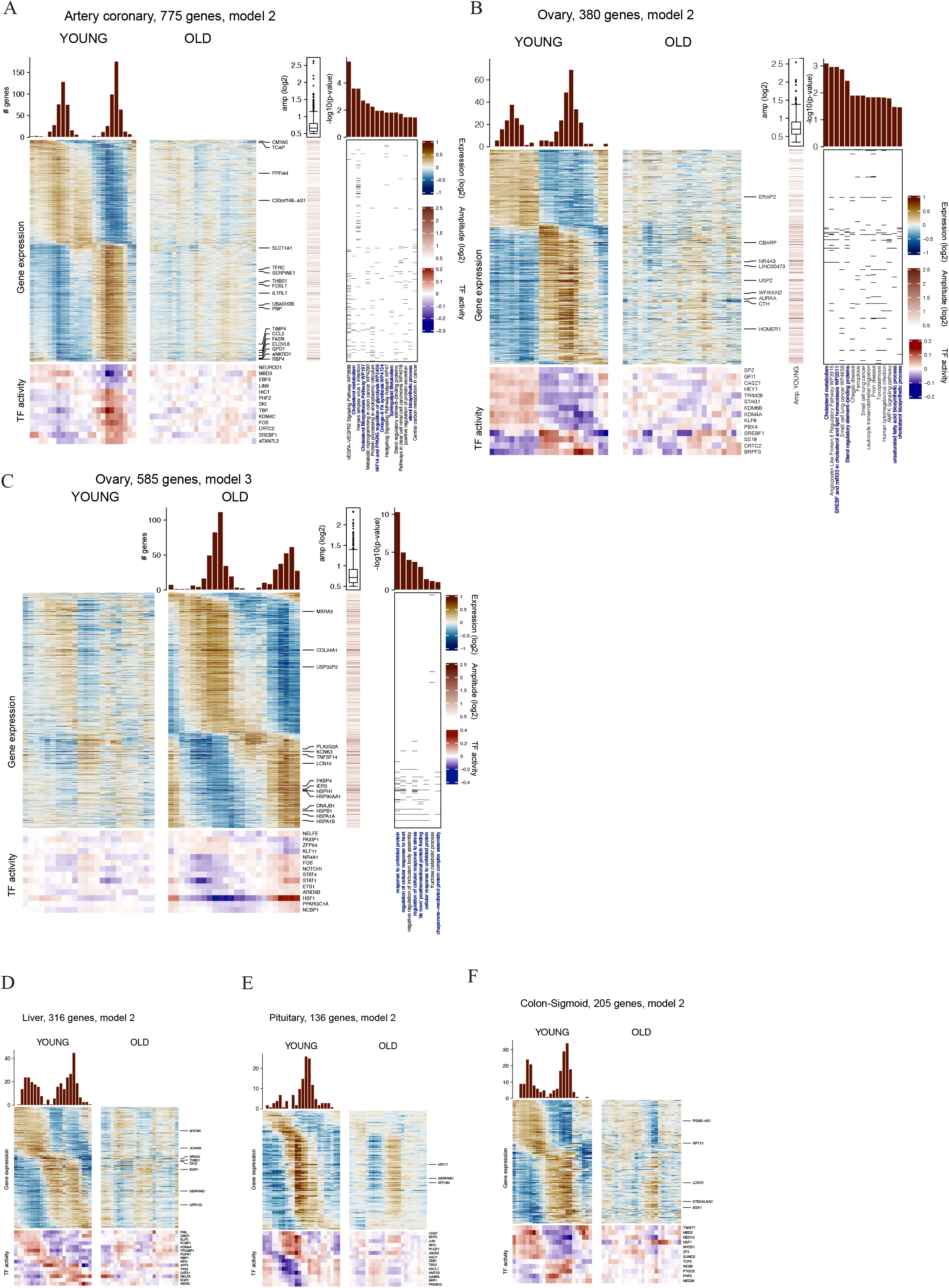
Compound heatmap visualizations of age dependent rhythmic mRNA programs in selected tissues. A) Artery coronary. Genes showing rhythmicity in younger donors analyzed and displayed as in Fig. S3A. Lipid, cholesterol and glucose metabolism terms are highlighted in blue text. B) Ovary gland. Genes only rhythmic in younger donors displayed as in A). Note the loss of rhythmic lipid metabolic functions in elderly. C) Ovary gland. Genes only rhythmic in older donors displayed as in A). Note the emergence of marked heat shock response rhythms in the older group. D) to F) Genes only rhythmic in young liver, pituitary and colon displayed as in A). Note that high amplitude 24h patterns in the younger group showed the emergence of 12h rhythms in older donors.

