## Supplemental method for "Sex-dimorphic and age-dependent organization of 24 hour gene expression rhythms in human"

June 13, 2022

### 1 The CHIRAL algorithm

Here we discuss the theory, mathematical steps, and approximations of our Circular HIerarchical Reconstruction ALgorithm, CHIRAL. This algorithm takes as input an ensemble of sets of measurements of quantities in a periodic system, like mRNA expression levels of core clock transcripts from tissue samples across the 24h. Each set of measurements represents a contemporary measurement of all the quantities in our system; In this example the set of measurements is the set of mRNA expression levels (RNA-Seq) for all clock genes for one sample. The aim of the algorithm is to assign to each set of samples in the ensemble a phase along the period of the system; i.e. a phase between 0 and 24hs for the circadian clock. We will keep in mind the circadian clock example throughout the calculations as that is our main practical objective.

#### 1.1 Multivariate harmonic regression model

As a starting point, we consider a number of genes by number of conditions (samples), $N_g \times N_c$ data matrix $E^g_c$. We still have in mind periodic behaviour, so the most intuitive basis in which to expand our signal is the Fourier basis. In particular we will consider up to $N_f$ nonzero components of the Fourier basis. With this in mind, we can write:

$$E^g_c = \alpha^g_f \zeta^f(\varphi_c) + \varepsilon^g_c = \alpha^g_f \zeta^f_c + \varepsilon^g_c \tag{1}$$

using the differential geometry notation where high indices only sum with low ones and vice versa. In this equation the $\alpha$s represent the gene coefficients associated with the Fourier functions $\zeta^f$ which are the same for all the possible phases $\varphi_c$. From (1) we can remind how the aim of the model is to obtain the phases $\{\varphi_c\}$ from the data $\{E_{gc}\}$. We use a compact notation for the Fourier series but see paragraph (1.2) for the explicit expression with phases. Here the error has two indices, as its distribution can depend both on the gene and on the condition. In addition we want a model that separates the genes between rhythmic genes and non rhythmic ones; the difference in this formulation between rhythmic and non-rhythmic lies in the prior over $\alpha$, as for non-rhythmic genes only the flat (DC or infinite period) Fourier harmonic is considered.

Before proceeding, let us specify the various probability distributions. We start with the distribution of random errors

$$\mathcal{P}(\varepsilon) \sim e^{-\frac{1}{2}\varepsilon_c^g S_{gg'}^{cc'} \varepsilon_{c'}^{g'}} \;, \tag{2}$$

where the covariance matrix, here written as a 4 dimensional tensor. The prior on the $\{\varphi\}$ is taken as flat. The probability on the $\{\alpha\}$ conditioned on the set of states $\{s\}$ will grow in complexity as we increase the number of Fourier components, as:

$$\mathcal{P}(\alpha|\{s\}) \sim e^{-\frac{1}{2}\alpha_f^g A(\{s\})_{gg'}^{ff'} \alpha_{f'}^{g'}}. \tag{3}$$

and the covariance matrix is written as $A$. We have introduced here $\{s\}$, the set of the state variables; for each gene the state indicates whether the gene is rhythmic, $s_g = 1$, or non rhythmic, $s_g = 0$. While these covariance matrices are general, we now make approximations on both.

#### 1.2 Approximations on the covariance matrices

Here we discuss approximations that will make calculations more tractable. We also drop the high-low indices notation for the easier Einstein notation where two same indices are summed ($a_i b^i$ becomes equal to $a_i b_i$). We start by simplifying the error covariance matrix:

$$(S^{-1})^{gg'}_{cc'} = \delta_{cc'}\delta^{gg'}\sigma^2 \ , \tag{4}$$

and then that of the probability for $\{\alpha\}$

$$(A(\{s\})^{-1})^{gg'}_{ff'} = \delta_{ff'}\delta^{gg'}\left(\delta_{0f}u^2 + \delta_{1s_g}\delta_{1\lfloor\frac{f}{2}\rfloor}\tau^2\right) \ , \tag{5}$$

which assumed a single Fourier mode. The model can then be rewritten as:

$$\begin{aligned} E_{gc} &= \nu_g + \varepsilon \quad \text{if s} = 0, \\ E_{gc} &= \mu_g + a_g\cos(\varphi_c) + b_g\sin(\varphi_c) + \varepsilon_{gc} = \alpha_g^\intercal\,\zeta_c + \varepsilon \quad \text{if s} = 1; \end{aligned} \tag{6}$$

with $\alpha_g^\intercal = (\mu_g, a_g, b_g)$, and $\zeta_c^\intercal = (1, \cos(\varphi_c), \sin(\varphi_c))$. As for the prior if $s = 1$:

$$\mathcal{P}(\alpha) = \sqrt{\frac{\det(T^{-1})}{(2\pi)^3}}e^{-\frac{1}{2}\alpha^\intercal T^{-1}\alpha}, \tag{7}$$

with

$$T = \begin{pmatrix} u^2 & 0 & 0 \\ 0 & \tau^2 & 0 \\ 0 & 0 & \tau^2 \end{pmatrix}. \tag{8}$$

and the error distribution:

$$\mathcal{P}(\varepsilon) \sim \prod_{gc} e^{-\frac{\varepsilon_{gc}^2}{2\sigma^2}} \ . \tag{9}$$

For the moment we will consider all genes to have $s = 1$ (one state model), and we will reintroduce the two state model with the Expectation-Maximization algorithm in paragraph(1.6). From $\mathcal{P}(\varepsilon)$ we can write:

$$\mathcal{P}(E|\{\varphi\}, \{\alpha\}) \sim \prod_{gc} e^{-\frac{(E_{gc}-\alpha_g\zeta_c)^2}{2\sigma^2}} = \prod_g \mathcal{P}(E_g|\{\varphi\}, \alpha_g). \tag{10}$$

After some simple algebra we obtain:

$$\mathcal{P}(E_g|\{\varphi\},\alpha_g)\mathcal{P}(\alpha_g) \sim e^{-\frac{1}{2\sigma^2}\left(\alpha_g^\intercal(M+\sigma^2T^{-1})\alpha_g\right)-2\alpha_g\cdot W_g}, \tag{11}$$

with

$$M = \sum_c \zeta_c^\intercal \zeta_c =: X^\intercal X, \quad W_g = \sum_c E_{gc}\zeta_c \ . \tag{12}$$

As the goal is to identify the sample phases $\varphi$ ($\zeta$s) without being interested in the $\alpha$'s, we integrate out the latter. After a multidimensional Gaussian integration we get:

$$\int \mathcal{P}(E_g|\{\varphi\},\alpha_g)\mathcal{P}(\alpha_g)d\alpha_g \sim e^{\frac{1}{2\sigma^2}W_g^\intercal(M+\sigma^2T^{-1})^{-1}W_g} \ . \tag{13}$$

To obtain the $\zeta$s, we rewrite this as:

$$\mathcal{P}(\varphi|E) \sim e^{\sum_{ij}\zeta_i K_{ij}\zeta_j} \sim e^{-\beta H[\varphi]} \tag{14}$$

with

$$K_{ij} = \sum_g E_{gi}M_\sigma^{-1}E_{gj} \quad M_\sigma = M + \sigma^2 T^{-1} \ . \tag{15}$$

Here, we introduced the physics notation where $\beta$ is the inverse temperature and $H$ a Hamiltonian. We remind that each $K_{ij}$ is a matrix, thus $K$ is a 4 dimensional tensor of the form $K_{ij}^{ab}$.

#### 1.3 Rewriting the problem as a spin model

For the moment we wrote the problem in function of $\zeta_c^\intercal = (1, \cos(\varphi_c), \sin(\varphi_c))$. However we can define a new variable that still carries all the phase information: $\eta_c^\intercal = (\cos(\varphi_c), \sin(\varphi_c))$. Doing this transformation will bring a more informative, although slightly more complicated, Hamiltonian. We define the $J_{ij}$ matrix as the bottom right $2 \times 2$ matrix of $K_{ij}$, i.e. $J_{ij}^{ab} = K_{i,j}^{a+1,b+1}$

with $a, b \in \{1, 2\}$, and the $h$ vector as the last two elements of the first column of $K$, i.e. $h_i^a = K_i^{a+1,3}$ with $a \in \{1, 2\}$. We notice how $K_{1,1}$ just contributes to a constant that will be included in the normalization of the probability which we are ignoring for the moment. So we have:

$$\mathcal{P}(\varphi|E) \sim e^{-\beta H[\varphi]} \tag{16}$$

with

$$H[\varphi] = -\left(\sum_{ij=1}^{N_c} \eta_i J_{ij} \eta_j + \sum_{i=1}^{N_c} h_i \eta_i\right), \quad \beta = 1 \tag{17}$$

with $J$ and $h$ defined above. Here we observe two canonical contributions to an XY spin model. The first term in the Hamiltonian is an all-to-all spin interaction term with couplings $J_ij$. Then we have a site-dependent external field term $h_i$ that attempts to align all spins to an external local field .

#### 1.4 Mean field approximation

Here we will derive a mean field approximation that allows us to calculate the unknown sample phases from a simple recursion relation. First, we take a closer look a the $M$ matrix, which can be written as:

$$M = \begin{pmatrix} 1 \cdot 1 & 1 \cdot C & 1 \cdot S \\ C \cdot 1 & C \cdot C & C \cdot S \\ S \cdot 1 & S \cdot C & S \cdot S \end{pmatrix} \quad A \cdot B = \sum_c A_c B_c \ , \tag{18}$$

with $C = (\cos(\varphi_1), \ldots, \cos(\varphi_{N_c}))$, $S = (\sin(\varphi_1), \ldots, \sin(\varphi_{N_c}))$.

The problem greatly simplifies when the samples are fairly uniformly distributed around the $24h$ cycle, which leads to the approximation that

$$C \cdot 1 = S \cdot 1 = C \cdot S = 0, \quad C \cdot C = S \cdot S = N_c/2 \tag{19}$$

Then, $M$, and also $M_\sigma$ become diagonal, and thus the above matrix inversion (Eq.15) will be much easier. Since $h_i = 0$ we obtain:

$$\mathcal{P}(\varphi|E) \sim e^{-\beta H[\varphi]} \tag{20}$$

with

$$H[\varphi] = -\sum_{ij} \eta_i J_{ij} \eta_j, \tag{21}$$

where

$$J_{ij} = \frac{1}{N_c N_g} \sum_g E_{gi} E_{gj}, \quad \beta = \frac{N_g}{\sigma^2} \frac{(N_c \tau^2)}{(\tau^2 N_c + 2\sigma^2)} \ . \tag{22}$$

We now derive a mean field approximation to obtain the expected phases contained in $\langle \eta_i \rangle$ (as $\eta_i = \eta_i(\varphi_i)$). We remind for clarity:

$$\mathcal{P}(\eta|E) \sim e^{\beta \sum_{ij} \eta_i J_{ij} \eta_j} \tag{23}$$

We start from the set of coupled equations:

$$\langle \eta_i \rangle = \frac{1}{Z} \int d\varphi_i e^{\beta \sum_j J_{ij} \eta_i \eta_j} \eta_i(\varphi) \tag{24}$$

which then reduces using the mean-field for $\eta_j$ in the exponent to:

$$\langle \eta_i \rangle = \frac{\int d\varphi_i e^{\beta \eta_i \sum_j J_{ij} \langle \eta_j \rangle} \eta_i}{\int d\varphi_i e^{\beta \eta_i \cdot \sum_j J_{ij} \langle \eta_j \rangle}}. \tag{25}$$

Since

$$\eta_i \sum_j J_{ij} \langle \eta_j \rangle = W_i \cos(\varphi_i - \bar{\varphi}_i)$$
$$\text{with} \quad \vec{W}_i \equiv W_i \hat{W}_i \equiv W_i \begin{pmatrix} \cos(\bar{\varphi}_i) \\ \sin(\bar{\varphi}_i) \end{pmatrix} := \sum_j J_{ij} \langle \eta_j \rangle \ , \tag{26}$$

where we have decomposed the vector $\vec{W}_i$ into the product of a unit vector $\hat{W}_i$ and a modulus $W_i$. Eq.(25) can be solved easily using Bessel functions, in particular:

$$\langle \eta_i \rangle = \frac{I_1(\beta W_i)}{I_0(\beta W_i)} \hat{W}_i \ . \tag{27}$$

This expression gives us a recursive method to calculate the phases. While this method is quick, its accuracy is limited owing to the many approximations made. However, the mean-field solution plays a crucial role as we use it to seed a much more accurate algorithm, which we now describe.

#### 1.5 The two-state model

While the phases inferred in by this XY mean-field model are typically ordered properly when tested in datasets with known time stamps (e.g. we used [22]), the actual values are not sufficiently accurate.

#### 1.6 The EM algorithm

To develop a more precise method to infer circadian phases we will proceed with an EM approach [61]. We will no longer assume that the genes are all in state one ($s = 1$) and consider a mixture model for states $s \in \{0, 1\}$, nor that the phases are well distributed around the $24h$. The rationale for the mixture model is to allow that some genes may not carry phase information in all the sample sets. In the iterative EM scheme, we will denote the new objects at any given step of the algorithm with a prime ($'$). We will also drop the gene index and the vector notation, as now the objects in play should be clear. The $Q$-function in the EM algorithm takes the form:

$$Q(\theta|\varphi) = \langle \log(\mathcal{P}(E, \alpha, s|\theta)) \rangle_{\mathcal{P}(\alpha,s|E,\varphi)} \ . \tag{28}$$

In particular, the phases at the next step are:

$$\varphi' = \arg\max_{\theta}(Q(\theta|\varphi)) \tag{29}$$

From now on we will abuse our notation and generally substitute $\theta$ with $\varphi'$ so that (28) becomes:

$$Q(\varphi'|\varphi) = \langle \log(\mathcal{P}(E, \alpha, s|\varphi')) \rangle_{\mathcal{P}(\alpha,s|E,\varphi)} \tag{30}$$

and (29)

$$\varphi' = \arg\max_{\varphi'}(Q(\varphi'|\varphi)) \ . \tag{31}$$

As our aim is to maximize $Q$ which involves taking a logarithm we can forget about some normalization and we can write from now on:

$$\mathcal{P}(E, \alpha, s|\varphi') = \mathcal{P}(E|\alpha, s, \varphi')\mathcal{P}(\alpha, s|\varphi') = \mathcal{P}(E|\alpha, s, \varphi')\mathcal{P}(\alpha|s)\mathcal{P}(s) \ . \tag{32}$$

Later in the calculation, we will need the probability that each gene is rhythmic ($s = 1$). To do so we need to apply Bayes theorem:

$$\mathcal{P}(s|E, \varphi) = \frac{\mathcal{P}(E|s, \varphi)\mathcal{P}(s)}{\sum_s \mathcal{P}(E|s, \varphi)\mathcal{P}(s)}, \tag{33}$$

where:

$$\mathcal{P}(E|s, \varphi) = \int d\alpha \mathcal{P}(E|\alpha, s, \varphi)\mathcal{P}(\alpha|s, \varphi), \tag{34}$$

and where we have used that the prior $\mathcal{P}(s|\varphi) = \mathcal{P}(s)$. Our prior on $\alpha$ is, depending on the state:

$$\mathcal{P}(\alpha_g|s = 1, \varphi) = \sqrt{\frac{\det(T^{-1})}{(2\pi)^3}} e^{-\frac{1}{2}\alpha_g^\intercal T^{-1}\alpha_g}. \tag{35}$$

or

$$\mathcal{P}(\alpha_g|s = 0, \varphi) = \delta(\alpha_g) \, . \tag{36}$$

As

$$\log(\mathcal{P}(E, \alpha|\varphi')) = \log(\prod_g \mathcal{P}(E, \alpha|\varphi')_g) = \sum_g \log(\mathcal{P}(E, \alpha|\varphi')_g) \tag{37}$$

taking into account (32), we have for each gene, considering only what we will need for the maximization:

$$Q(\varphi'|\varphi) = \int d\alpha\, (E - X'\alpha)^2 e^{\frac{1}{2\sigma^2}(\alpha-\hat{\alpha})^\intercal M_\sigma(\alpha-\hat{\alpha})}\mathcal{P}(s = s^1|E, \varphi) \tag{38}$$

where $X'$ his the same as implicitly defined in (12) and we define $t_g := \mathcal{P}(s_g = s^1|E, \varphi)$.

Solving the integral and putting everything together yields for the part that will be relevant for the maximization of $Q$: :

$$Q(\varphi'|\varphi) = -\sum_g t_g(E_g^\intercal E_g - 2E_g^\intercal X'\hat{\alpha_g} + \hat{\alpha_g} M'\hat{\alpha_g}) - \mathrm{Tr}(\sigma^2 M_\sigma^{-1} M') \tag{39}$$

So all that we need to still do is calculate all the $t_g$ weights. Before we begin the calculation, we need to correctly normalize all the probabilities in play:

$$\int \mathcal{P}(E|\alpha, s^1, \varphi) \prod dE = \int e^{-\frac{(E-\alpha\zeta)^2}{2\sigma_1^2}} dE = (2\pi\sigma_1^2)^{M/2} = Z_E^1 \tag{40}$$

with $M$ the number of entries of the $E$ matrix. For state zero ($s = 0$) we find

$$\int \mathcal{P}(E|\alpha, s = 0, \varphi) \prod dE = \int e^{-\frac{E^2}{2\sigma_0^2}} dE = (2\pi\sigma_0^2)^{M/2} = Z_E^0. \tag{41}$$

We now do all the calculations omitting the gene index, as it only burdens the notation. Marginalizing the $\alpha$ yields

$$\begin{aligned}
\mathcal{P}(E|s = 1, \varphi) &= \int d\alpha \mathcal{P}(E|\alpha, s = 1, \varphi)\mathcal{P}(\alpha|s = 1, \varphi) && (42)\\
&= \frac{1}{Z_E^1}\sqrt{\frac{\det(T^{-1})}{\det(M_\sigma/\sigma_1^2)}} e^{\frac{1}{2\sigma_1^2}(E^\intercal X M_\sigma^{-1} X^\intercal E - E^\intercal E)} . && (43)
\end{aligned}$$

In addition, we can also easily calculate the other probability ($s = 0$):

$$\mathcal{P}(E|s = 0, \varphi) = \frac{1}{Z_E^0} e^{-\frac{E^\intercal E}{2\sigma_0^2}} . \tag{44}$$

Finally we obtain the gene weight

$$t = \frac{\gamma q e^{\frac{1}{2\sigma^2}(E^\intercal X M_\sigma^{-1} X^\intercal E)}}{1 - q + \gamma q e^{\frac{1}{2\sigma^2}(E^\intercal X M_\sigma^{-1} X^\intercal E)}} \ , \tag{45}$$

with

$$\gamma = \left(\frac{\sigma_0^2}{\sigma_1^2}\right)^{M/2} \sqrt{\frac{\det(T^{-1})}{\det(M_\sigma/\sigma^2)}} e^{\frac{E^\intercal E}{2}\left(\frac{1}{\sigma_0^2} - \frac{1}{\sigma_1^2}\right)} \ . \tag{46}$$

Note that in these calculations we have mean-centred our data matrix $E$. However, this does not mean that $\mu_g$ is zero in this model (Eq. 6) since the density of phases may not be constant along the cycle.

The final expression for $Q$ and its derivatives reads:

$$Q(\varphi'|\varphi) = -\sum_g t_g \left( E_g^\intercal E_g - 2E_g^\intercal X' \hat{\alpha}_g + \hat{\alpha}_g M' \hat{\alpha}_g - \text{Tr}({\sigma_g}^2 M_{\sigma_g}^{-1} M') \right) \tag{47}$$

$$0 = \frac{\partial Q(\zeta'|\zeta)}{\partial C_i'} =$$

$$\sum_g t_g \left( \hat{a}_g(-E_{gi} + \hat{\mu}_g + \hat{a}_g C_i + \hat{b}_g S_i) + (M_\sigma^{-1})_{2,1} + (M_\sigma^{-1})_{2,2} C_i + (M_\sigma^{-1})_{2,3} S_i \right) = 0 \tag{48}$$

$$0 = \frac{\partial Q(\zeta'|\zeta)}{\partial S_i'}$$

$$\sum_g t_g \left( \hat{b}_g(-E_{gi} + \hat{\mu}_g + \hat{a}_g C_i + \hat{b}_g S_i) + (M_\sigma^{-1})_{3,1} + (M_\sigma^{-1})_{3,2} C_i + (M_\sigma^{-1})_{3,3} S_i \right) = 0. \tag{49}$$

This set of equation can be compactly rewritten as:

$$K \overline{\zeta}_i = \overline{O} \tag{50}$$

with the evident definitions.

However, since we want to keep the constraint that $|\zeta|^2 = 1$ we need to introduce Lagrange multipliers. Reminder: in general, we want to maximize $f(\overline{x})$ where $\overline{x}$ satisfies $g(\overline{x}) = 0$. To do this, we can maximize $h(\overline{x}, \lambda) =$

$f(\overline{x}) + \lambda g(\overline{x})$ (with respect to all its variables). If there is more than one constraint, we need to introduce more than one multiplier: one for each constraint. In our case:

$$h(\zeta', \lambda) = Q(\zeta'|\zeta) + \sum_i \lambda_i (C_i^2 + S_i^2 - 1). \tag{51}$$

So, as before, if we consider sample $i$ we have:

$$(K + \lambda I)\overline{\zeta}_i = \overline{O} \tag{52}$$

and

$$C_i^2 + S_i^2 = 1. \tag{53}$$

So we need to find the $\lambda$ such that

$$\left|(K + \lambda I)^{-1}\overline{O}\right|^2 = 1 \ . \tag{54}$$

To solve (54), and without a better solution, we calculate explicitly the fourth order polynomial in $\lambda$ of which the roots are the maxima or minima. As anyone would easily expect, what comes out is indeed very ugly. If we write:

$$\overline{O} = \begin{pmatrix} \alpha \\ \beta \end{pmatrix} \qquad K = \begin{pmatrix} A & B \\ C & D \end{pmatrix} \ , \tag{55}$$

we find

$$\begin{aligned} &\lambda^4 + \lambda^3(2A + 2D) + \lambda^2(D^2 + A^2 + 4AD - \alpha^2 - \beta^2 - 2BC) + \\ &+ \lambda(2AD(A + D) - 2BC(A + D) - 2D\alpha^2 - 2A\beta^2 + 2\alpha\beta(B + C)) + \\ &+ B^2C^2 + A^2D^2 - \alpha^2(D^2 + C^2) - \beta^2(A^2 + B^2) + 2\alpha\beta(AB + CD) - 2ABCD = 0. \end{aligned} \tag{56}$$

Solving this numerically provides us with up to 4 different possible values of $\varphi'$, so in order to find which gives rise to the max of $Q$ we simply evaluate the function in all the 4 points. This can be done separately for each condition as the derivatives become independent from one another. With this we have solved the minimization process and have found our $\varphi'$ thus we have concluded an iteration of the EM.

#### 1.7 Practicalities

Up to now we have considered the set of all genes in the given set of mRNA measurements. However, given the completely unsupervised nature of this algorithm we cannot, a priori, be sure of which oscillatory process will be picked up (e.g. cell cycle or circadian clock). To be sure that we are capturing the circadian time as a source of variation we can apply our algorithm to a subset of genes. In particular, in this case, we selected the 12 clock reference genes (CRGs): DBP, PER3, TEF, NR1D2, PER1, PER2, NPAS2, ARNTL, NR1D1, CRY1, CRY2, CIART. Also, we found that the prior on the $\alpha$s needs to be tightly controlled as it "competes" against the experimental data and we know that genes do not have oscillation of more than $10 - 20$ log2 units. So for the prior we pick $u^2 = 0.2$, $\tau^2 = 4/(24 + n)$ where $n$ is the number of samples.
